# Pigments and microstructure of the colour polymorphic shells of *Polymita picta* and *P. muscarum* (Gastropoda: Cepolidae), with observations on a new light-transmitting shell spot system

**DOI:** 10.64898/2026.03.03.709309

**Authors:** Mario Juan Gordillo-Pérez, Natalie Beenaerts, Julia Sigwart, Thierry Backeljau, Thomas Vranken, Javier Vilasó-Cadre, Martijn Heleven, Karen Smeets, Dries Vandamme, Bernardo Reyes-Tur

**Affiliations:** Hasselt University, Centre for Environmental Sciences, Hasselt, Belgium; Senckenberg Research Institute and Museum of Natural History Frankfurt, Frankfurt, Germany; Royal Belgian Institute of Natural Sciences, OD Taxonomy and Phylogeny, Brussels, Belgium; University of Antwerp, Department of Biology, Antwerp, Belgium; Hasselt University, Institute for Materials Research (IUMAT), Hasselt, Belgium; imec, IUMAT, Diepenbeek, Belgium; EnergyVille, IUMAT, Genk, Belgium; Universidad del Valle de México, San Luis Potosí, México; University of Oriente, Department of Biology and Geography, Santiago de Cuba, Cuba

## Abstract

Colour polymorphism in the Cuban painted snails *Polymita picta* and *P. muscarum* is striking, yet the pigmentary and structural bases remain unclear. We combined spectrophotometric screening, Raman micro-spectroscopy, scanning electron microscopy (SEM) and LED transillumination to link pigments, ultrastructure and optics across shell morphs. A *Sepia officinalis* melanin standard yielded a robust linear calibration used to quantify total melanin pigments at 215 nm in pooled extracts. Melanin was detected in all samples with predominance in darker morphs. Raman spectra (785 nm) confirmed aragonite mineral organization and revealed carotenoid bands, consistent with a mixed-pigment model in which carotenoids contribute to ground and band colours and melanins underlie darker elements. SEM showed a canonical crossed-lamellar layer with alternating transverse and co-marginal layers. At shell spot areas, shell surfaces were cribose; fracturing shells near such spots showed a locally disordered, more porous mineral arrangement enriched in organic matrix, bounded basally by an organic layer. Under transillumination, these spots acted as discrete light-transmitting windows, abundantly in *P. muscarum* and also sparsely in *P. picta*; as the spots appear only as pigmented dots but also transmit light, we refer to them as ‘cryptotransmissive’ domains, which is apparently a novel finding in terrestrial gastropods. We speculate that these may represent a pigment-structure-optics framework in which pigments and microstructural packing jointly play potential roles in photoprotection and behavioural thermoregulation. These results provide a mechanistic context for colour polymorphism in *Polymita* and suggest testable links to thermal ecology and conservation.

## INTRODUCTION

Mollusc shells exhibit an extraordinary diversity of colours, patterns and degrees of transparency resulting from the combined effects of pigment chemistry and shell ultrastructural organization. Together, these factors generate a wide range of optical behaviours, including absorption, scattering and transmission of light. Shell colour has long attracted attention of researchers due to its ecological significance, as it plays roles in camouflage, signalling, photoprotection, thermoregulation, and immune defence (Scheil et al., 2013; Mann and Jackson, 2014; Williams, 2017; Davison et al., 2019; Checa et al., 2018; Stuart-Fox et al., 2021; Gefaell et al., 2023). As such, shell colour provides a useful system for studying the evolutionary meaning and dynamics of phenotypic polymorphisms. Despite decades of research, major knowledge gaps remain regarding the chemical identity of shell pigments, the structural organization of the pigments in the crystalline architecture of the shell, and the functional role(s) of shell colour traits in natural populations (Cuthill et al., 2017; Williams, 2017).

Three main classes of pigments are commonly associated with molluscan shell colour: melanins, carotenoids and porphyrins (Hedegaard et al., 2006; Williams, 2017). Melanins are typically linked to darker patterns and bands, carotenoids contribute to yellow, orange and pink hues, and porphyrins have been associated with a variety of shell colours which may fluoresce under UV light (Speiser et al., 2014; Williams, 2017). However, these associations are not universal, and pigment composition can vary considerably even among closely related taxa (Verdes et al., 2015; Affenzeller et al., 2020). Moreover, research on molluscan shell pigments studies has predominantly focus on marine species (e.g. Affenzeller et al., 2019b; Hu et al., 2025)

The genus *Polymita* H. Beck, 1837 (Gastropoda: Cepolidae) comprises six species of tree snails endemic to Eastern Cuba (Lewis et al., 2025) and the group is renowned for its striking shell colour polymorphism (Reyes-Tur et al., 2025). Yet, the shell pigments that are responsible for shell colour in *Polymita* were hitherto only investigated by Comfort (1951), who focussed on *Polymita picta* (Born, 1778). That study reported the presence of melanins, no detectable amounts of purine-based pigments, and none of the shell colour bands exhibited UV fluorescence, suggesting that porphyrin-type pigments (e.g. uroporphyrins) are likely absent or present at undetectable levels. To date, no comprehensive analyses have assessed pigment diversity, relative abundance or spatial distribution within *Polymita* shells, severely limiting our understanding of the molecular and evolutionary bases of the conspicuous colour polymorphism in *Polymita*.

Shell colour can have important functional consequences by influencing thermoregulation, providing photoprotection, and interacting with predators and parasites (Stuart-Fox et al., 2021; Gordillo-Pérez et al., 2025). Such interactions may contribute to the dynamics of colour polymorphism through selective pressures, including frequency-dependent predation and parasite-mediated selection (Reid, 1987; Johannesson & Ekendahl, 2002; Rosin et al., 2011; Surmacki et al., 2013; Mendonça et al., 2014). In taxa such as *Polymita* that can inhabit open, sun-exposed environments (González-Guillén, 2021), variation in pigment composition and shell ultrastructure may strongly affect heat absorption, thermal performance and tolerance to thermal stress (Seuront et al., 2018; Schweizer et al., 2019). In addition, highly conspicuous colour patterns may influence predator–prey interactions, suggesting that shell colour polymorphism could be maintained by multiple and potentially interacting selective pressures (Cuthill et al., 2017).

Gastropod shells are composite biominerals composed mainly of crystalline aragonite and/or calcite (both calcium carbonate) embedded within a minor, but functionally important, organic matrix. Externally, shells are covered by the periostracum, a chitin and protein-rich layer that templates early shell mineralization. Both shell and periostracum may contain pigments. At the microscopic scales, shells frequently exhibit a crossed-lamellar crystalline microstructure organized hierarchically into first-, second- and third-order lamellae, often combined with prismatic or other crystalline mineral structures (de Paula and Silveira, 2009; Song et al., 2019; Höche et al., 2020). These structural features can directly influence optical performance, since light propagation in calcium carbonate-based composites depends not only on pigment absorption, but also on packing density, porosity and refractive-index contrasts at organic-mineral interfaces (McCoy et al., 2024). The molluscan shell mineral arrangement can show various structural modifications, such as asymmetry in the prismatic layer and the formation of tubules, tunnels, or pores that extend fully or partially through the shell wall (e.g. Oberling, 1955, 1964; Batten, 1984; Kiel, 2004; Okada et al., 2009; Chen et al., 2021; Páll-Gergely, 2025). These structures have been associated with muscle insertions and/or the exchange of gases, water and ions.

Some shells include transparent light-transmitting domains. In some photosymbiotic bivalves, specialized aragonitic “windows” channel light in a manner similar to fibre-optic systems (McCoy et al., 2024). In some species of polyplacophorans (chitons), there are shell eyes with a crystal lens that is made of aragonite that focus light on photoreceptors embedded in the shell (Boyle, 1969; Li et al., 2015; Friedrich et al., 2021; Sigwart et al., 2025). Whether comparable light-transmitting mechanisms occur in terrestrial gastropods remains unexplored.

*Polymita picta* and *P. muscarum* (I. Lea, 1834) display the most striking shell colour diversity in this genus, including highly variable background colours and banding patterns. The shells also include irregularly distributed “spots” that generate diffuse black, brown or reddish areas in pigmented shells (de la Torre, 1950) and appear translucent in entirely white shells. These spots likely play a functional role in luminance modulation (Gordillo-Pérez et al., 2025). Although this extreme shell colour polymorphism has made *Polymita* a potential model for investigating the evolution and maintenance of shell colour diversity in terrestrial gastropods -particularly in Neotropical regions where such studies are scarce-the mechanisms underlying this variability remain poorly understood (Davison, 2002).

The six *Polymita* species in Cuba are threatened by climate change, habitat loss and poaching (CITES, 2025). As shell colour is one of the most conspicuous traits driving the exploitation of *Polymita* sp., elucidating the pigmentary and structural bases of shell colour is not only essential for evolutionary biology, but also highly relevant for conservation. Knowledge of the mechanisms underlying colour variation may help assess the loss of phenotypic diversity, identify potentially vulnerable morphs and evaluate the adaptive significance of colour under climate change. Therefore, in this study we investigate the pigment composition and shell ultrastructure across colour morphs of *P. picta* and *P. muscarum*, considering their potential ecological significance and implications for conservation.

## MATERIAL AND METHODS

### Samples and phenotype classification

Due to the threatened status of *Polymita picta* and *P. muscarum*, destructive sampling done was kept to a minimun. Whenever possible, visually well-preserved but previously broken shells belonging to the collection of the Terrestrial Molluscs Captivity Breeding Laboratory of Universidad de Oriente, Cuba were used to analyse pigments and shell ultrastructure. Shells of *Polymita picta* and *P. muscarum* were classified following established descriptions of shell colour polymorphism (Alfonso and Fernández, 1992; Alfonso and Berovides, 1989). In both species, shell colour variation arises from combinations of background (base) colour and the presence, number and colour of spiral bands.

In *P. muscarum*, two main base colours (white and brown) can be distinguished, which, in combination with banding patterns, give rise to multiple phenotypes. In *P. picta*, six base colours (white, yellow, red/orange, green, brown and black) combine with bands of varying colour (e.g. dark, pink or white) to generate a high diversity of phenotypes. Given this pronounced polymorphism, and for the purposes of the present study, shell regions were described operationally according to their visual characteristics (e.g. base colour, dark bands, pink bands), rather than using detailed morphotype codes. Additionally, entirely white shells of *P. muscarum* which have been referred to as var. *albina* or *albida* (González-Guillén, 2021), were included. This designation is used descriptively and does not imply genetically confirmed albinism. No permits were required for the described study, which complied with all relevant regulations.

### Melanin detection in shells using UV-visible spectrometry

Shells were pretreated and total melanin was extracted following Hao et al. (2015) and Affenzeller et al. (2020). Shell fragments were cut so as to obtain pieces with either only the background colours or only colour bands. These pieces were washed in deionized water, dried and weighed. Between 53.0 and 890.6 mg of shell belonging to 1–3 individuals per colour morph were placed in tubes with 3 mL of 6 mol/L HCl for over 12 h to hydrolyse shell proteins (Supplementary Data 1: https://doi.org/10.5281/zenodo.19185838). Next, these tubes were centrifuged at 3000 rpm for 5 min after which the supernatant was discarded. At this point, another 3 mL of 6 mol/L HCl was added to the tubes which were subsequently incubated for 1 h in boiling water to finish protein hydrolysis. The remaining undissolved portions were washed three times with deionized water and again centrifuged at 3000 rpm for 5 min. Next, the precipitated pigmented powder in each tube was dried at 80 °C and dissolved in 2 mL of 0.01 mol/L NaOH. For each precipitate, the UV absorbance spectrum (range of 190–500 nm) was scanned using a Cary 5000 UV-Vis-NIR Spectrophotometer from Agilent Technologies. The 0.01 mol/L NaOH solution was used as a blank. The presence of melanin was inferred from a broadband, monotonically decreasing absorbance from the UV to the visible range (Hao et al., 2015). Finally, curves were plotted using Cary WinUV software version 5.3.

### Melanin quantification in shells

A standard stock solution of commercial *Sepia officinalis* melanin (95%; Sigma-Aldrich, St. Louis, MO, USA) at 1 mg/mL was prepared in 0.01 mol/L NaOH under constant stirring for 2 h at room temperature to ensure complete dissolution in the dark. Working solutions were prepared by serial dilution of the stock to final concentrations of 40, 80, 160, 240, 320, 400 and 480 µg/mL. A solvent blank was measured at the beginning of each analytical run and used for baseline correction. Each concentration standard was prepared and measured in triplicate.

For descriptive purposes, the mean absorbance at 215 nm (Hao et al., 2015) and standard deviation (SD) were calculated for each concentration level. The calibration curve, however, was fitted in R (version 4.4.0) using ordinary least squares (OLS) linear regression on all individual replicate measurements (absorbance vs. concentration) (Supplementary Data 2: https://doi.org/10.5281/zenodo.19186051). Model quality was assessed using the coefficient of determination (*R²*), analysis of variance of the regression (F statistic), and residual diagnostics, including tests of normality and homoscedasticity. Analytical sensitivity parameters were derived from the regression output. The limit of detection (LOD) was calculated as LOD = 3σ/*S*, and the limit of quantification (LOQ) as LOQ = 3.33 × LOD (approximately 10σ/*S*), where *S* is the slope of the calibration line and σ is the standard error of the intercept obtained from the OLS fit.

For each shell pigment precipitate showing a melanin-like spectrum, melanin was quantified at 215 nm using the calibration based on commercial *Sepia officinalis* melanin. For each melanin-positive precipitate, melanin concentration (*C*) was calculated as *C* (µg/mL) = (*A*_(215)_ – *a*)/*S*, where *A*_(215)_ is the blank-corrected absorbance at 215 nm, and *a* and *S* are the intercept and slope of the calibration curve, respectively. Melanin mass (*M*) in the precipitate was calculated as *M*(µg) = *C* (µg/mL) × *Vf* (mL) × *DF*, with *Vf* = 2 mL (final volume of dissolved precipitate for all samples) and *DF* the corresponding dilution factor of the calibration solution (here 1). The amount of melanin relative to the processed shell mass was expressed as percentage (Lehmann et al., 1996), (Supplementary Data 1: https://doi.org/10.5281/zenodo.19185838).

Estimates outside the validated calibration range (40–480 µg/mL) were flagged as “extrapolated” and interpreted cautiously. Results below the LOD were reported as “melanin not detected”, whereas values between the LOD and LOQ were reported as “melanin detected but not reliably quantifiable”. Reported melanin percentages refer to pooled material from up to three individual shells per sample and therefore do not capture individual-level variability. The construction and evaluation of the calibration curve followed the criteria of Greenwell et al. (2014), Ranke et al. (2018), Conti (2022) and Chand et al. (2023).

### Raman spectroscopy

The shell base colour, as well as the colour of the dark, pink and white bands or spots on the surface of the shells were examined using microscopic analysis combined with Raman spectroscopy (Renishaw inVia Qontor) using a 785 nm, 100 mW laser. A 50x Leica long working distance objective was used, in combination with a 1200 lines/mm grating. Shell fragments were subjected to 10% laser power (∼ 10 mW), with an exposure time of 2 s per accumulation using 10 accumulations per spectrum. All measurements were performed under ambient conditions.

### Scanning electron microscopy (SEM) of shell ultrastructure

The preparation of *Polymita* shell fragments for SEM followed Carter and Ambrose (1989) and de Paula and Silveira (2009). Un-polished and un-etched shell fragments were washed by sonication in distilled water, dried and mounted on SEM stubs using double-sided conductive carbon adhesive tape. One replicate of each sample (a fragment of the same shell and region) was coated by gold sputtering to contrast with uncoated pieces. Three shell fragments containing spot areas from *P. muscarum* were hand-polished (not embedded) using silicon carbide abrasive papers (following FEPA P-grit abrasive particle size standards) under minimal pressure. These fragments originated from three different individuals. Due to limited material availability, spot areas were only analyzed in *P. muscarum*, and not in *P. picta*. Wet grinding was performed using distilled water with ∼30 % ethanol to limit carbonate dissolution and samples were frequently rinsed. The grit sequence was P320, P600, P1200, P2500 and P4000. The polishing direction was periodically rotated over ∼ 90 degrees and continued until scratches from the preceding grid were removed. Shell fragment surfaces were inspected under a stereomicroscope to confirm their flatness and the absence of deep scratches. The fragments were observed and photographed using a Phenom ProX scanning electron microscope (SEM) operated at 15 kV under low-vacuum conditions (1 Pa). Energy dispersive spectroscopy (EDS) was used to discriminate atomic compositions between mineral and organic matrix components in the shells. We proceeded to describe the shell microstructures following the terminology of Carter and Ambrose (1989), Chateigner et al. (2000) and de Paula and Silveira (2009). Because the ultrastructure of the shell spots suggested potential light-transmitting effects, shells were transilluminated by white light using an RGB LED base light (3-channel RGB SMD LEDs, Tangle Hardware and Plastic Products). For each trial, shells were placed over a diffuser plate, 2 cm from the light source and imaged inside a darkened enclosure to minimize stray light.

## RESULTS

### Melanin calibration curve, detection and quantification in shell fragments

A standard calibration curve was generated using commercial *Sepia officinalis* melanin. Absorbance values increased linearly with melanin concentration over the range of 40–480 µg/mL (Fig. 1). OLS regression yielded the equation *y* = 0.1606 + 0.0027*x*, with a coefficient of determination *R*² = 0.9907. The slope (*S* = 0.00266 ± 0.00006) and intercept (*a* = 0.1606 ± 0.0170) indicated high analytical sensitivity and low background signal, respectively. Residual diagnostics showed no significant deviation from normality (Shapiro–Wilk test, *p* = 0.451) and no evidence of heteroscedasticity (Breusch–Pagan test, *p* = 0.386), while the low residual standard error (0.041) confirmed the robustness of the model. The analysis of variance of the regression (F = 2028.6, *p* < 2.2 × 10^−16^) further supported its statistical significance. Based on the standard error of the intercept, the LOD was estimated at 19.2 µg/mL and the LOQ at 64.1 µg/mL. Together, these parameters demonstrate that the calibration curve is reliable and suitable for quantifying melanin in subsequent experimental samples.

**Figure 1.**
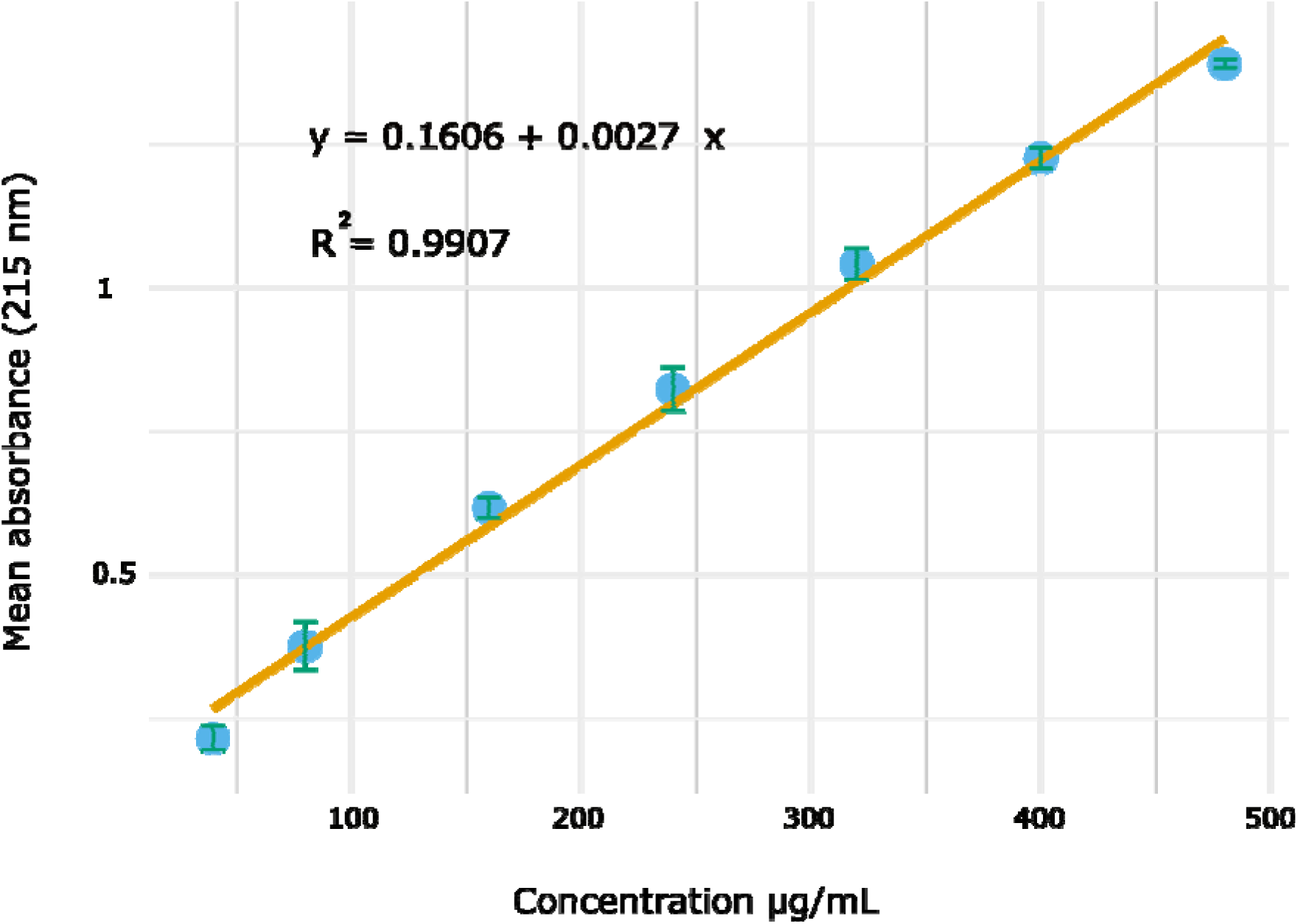
Calibration curve of the mean absorbances and their SD of the total melanin concentration in a standard commercial *Sepia officinalis* melanin dilution series of 40, 80, 160, 240, 320, 400 and 480 µg/mL at 215 nm.

Melanin was detected in all 15 shell-derived precipitates and quantified at 215 nm using the external standard calibration curve (Fig. 2), (Supplementary Data 1: https://doi.org/10.5281/zenodo.19185838). Nine of these extracts had melanin concentrations within the validated calibration range (40–480 µg/mL) and were classified as “in range”, whereas six exceeded this range and were classified as “extrapolated”. Values within the calibration range are considered quantitatively reliable. In contrast, extrapolated values (i.e. > 480 µg/mL) fall outside the calibration range and are therefore increasingly uncertain. Accordingly, extrapolated values are only used for qualitative comparisons (e.g. ranking of morphs) and not for precise concentration estimations. The nine extracts “in range” showed melanin concentrations above the LOQ (64.1 µg/mL), spanning 66.5–407.5 µg/mL (0.029–0.943%), with the highest concentration (0.293%) recorded in the dark band of the green morph of *P. picta*. The highest “extrapolated” melanin concentrations were observed in the black morph (0.944%), in the base colour of the red morph (0.553%), and in the dark band of the red morph (0.337%) of *P. picta*. At the lower end of the spectrum, the lowest concentrations were observed in the base colour of the brown morph (0.029%) of *P. picta*, followed by the base colour of brown morph (0.050%) and in the entirely white morph (0.059%) of *P. muscarum*.

**Figure 2.**
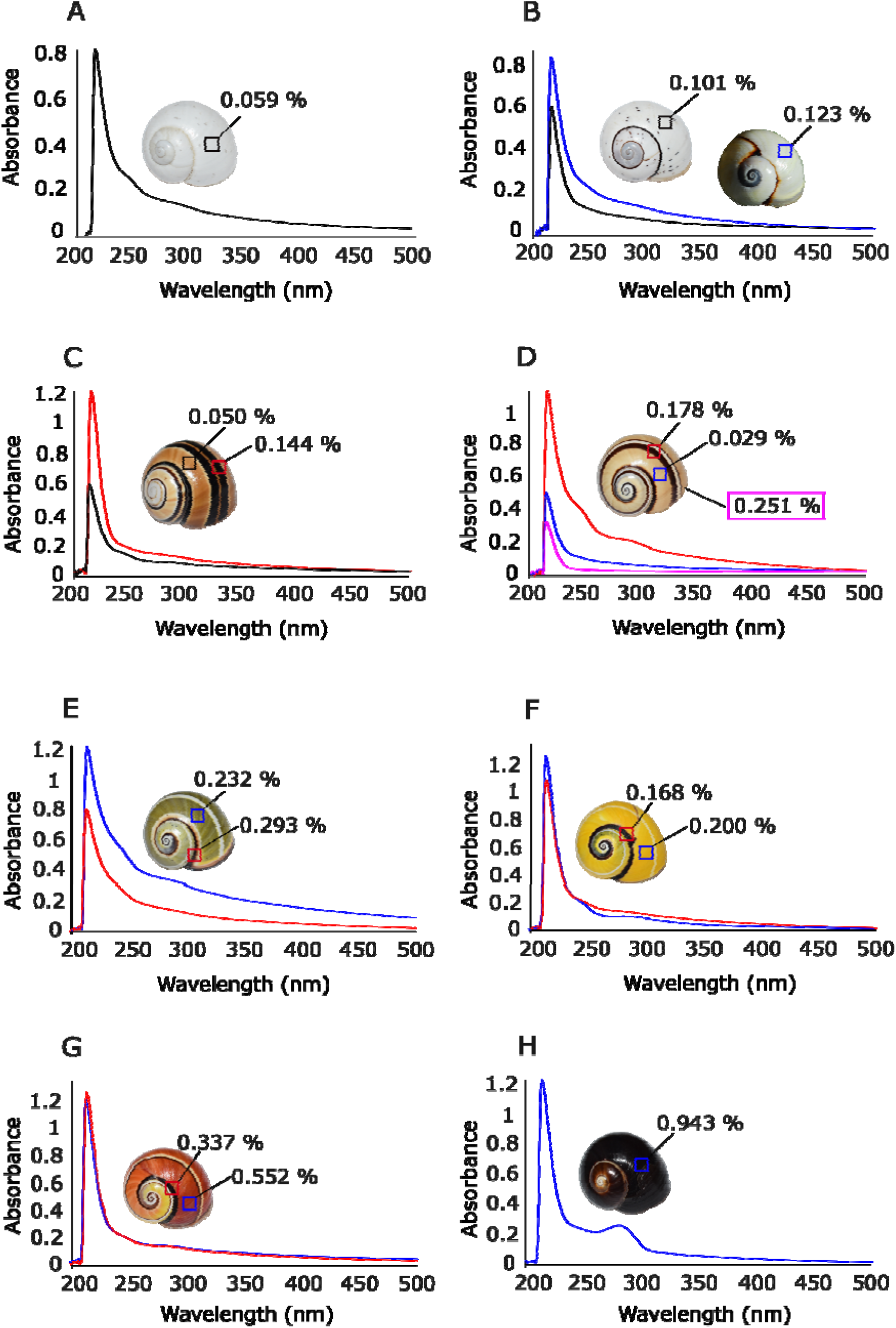
Melanin detection and quantification based on UV absorption spectra in different colour phenotypes of *Polymita picta* and *P. muscarum*. Melanin percentages are expressed relative to the mass of the pooled shell fragments of each sample (material from 1–3 individuals per sample). The shell areas analyzed are indicated by squares. A: *P. muscarum*, base colour of entirely white morph. B: *P. muscarum* and *P. picta*, base colour of white morph. C: *P. muscarum*, dark band and base colour of brown morph. D: *P. picta*, dark band, pink band and base colour of brown morph. E: *P. picta*, dark band and base colour of green morph. F: *P. picta*, dark band and base colour of yellow morph. G: *P. picta*, dark band and base colour of red-orange morph. H: *P. picta*, base colour of black morph.

### Raman spectroscopy

In *Polymita picta*, Raman spectra acquired with 785 nm excitation show a consistent mineral signature across all morphs, characterized by a strong peak at ∼1084 cm□¹ and a well-resolved double peak at 700/704 cm□¹ (Fig. 3). These peaks correspond to the ν□ and lattice modes of aragonite. No differences in mineral phase were detected among the examined colour morphs. In addition, several organic-compound-related bands are detected. In the base colour of the black morph, bands occur at approximately 1010, 1115, 1292 and 1494 cm^−1^, the latter appearing as a low-intensity shoulder. The dark band of the red morph exhibits the same set of peaks except for the 1292 cm^−1^ band. In the base colour of the brown morph, these pigment-related peaks are present but with much lower intensity. The base colour of the red morph displays well-defined bands at 1124 and 1509 cm^−1^, while the base colour in yellow and green morphs shows only low-intensity bands at 1136 and 1541 cm^−1^. Overall, darker morphs show stronger Raman signals in the 1100–1550 cm^−1^ region, consistent with higher organic pigment concentrations, whereas lighter morphs show weaker or nearly flat spectra.

**Figure 3.**
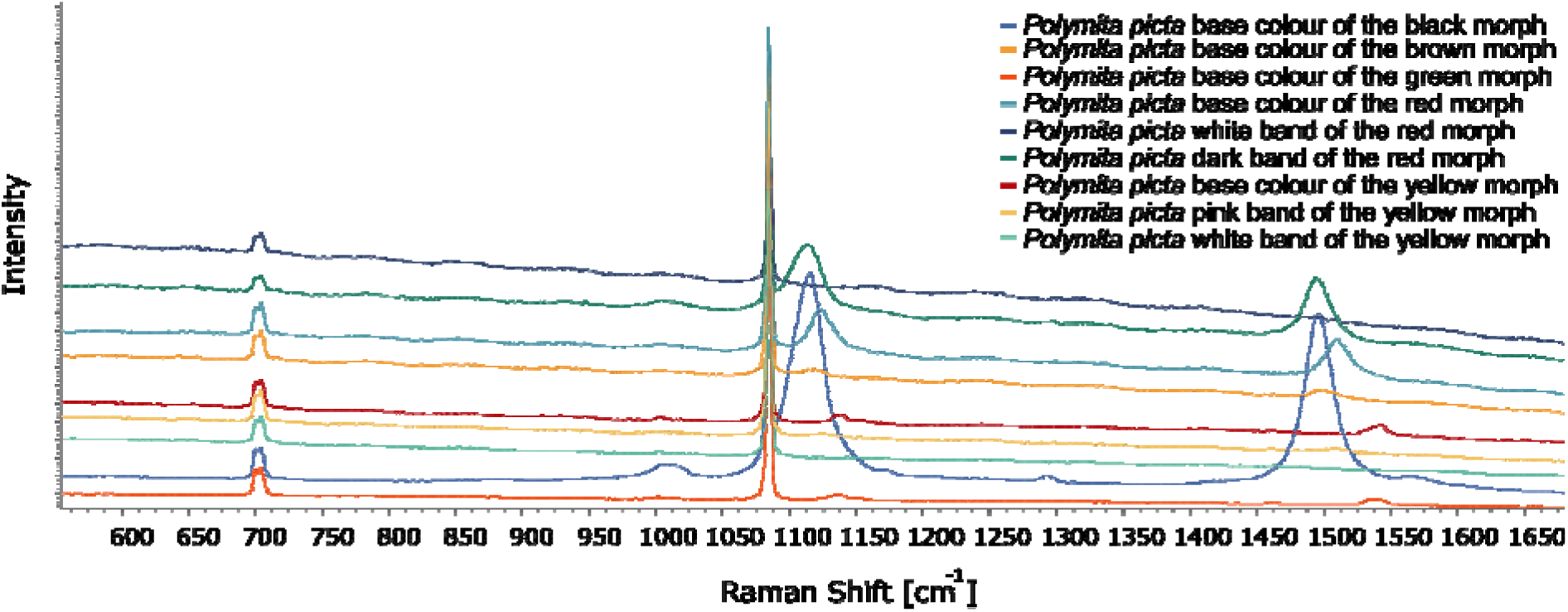
Raman spectra obtained from the base colour and bands in shell fragments of *Polymita picta*.

*Polymita muscarum* shows the aragonite signature of *P. picta*, with a strong peak at 1084 cm^−1^ and a well-resolved double peak at 700/704 cm^−1^ (Fig. 4). In the dark band of the brown morph, peaks occur at 1008, 1113, 1293 and 1493 cm^−1^, the latter presenting a noticeable shoulder. In the base colour of the brown morph and in the black spots of the white morph, well-defined bands at 1113 and 1493 cm^−1^ are observed. The spectral intensity of these bands is higher in darker shell areas, indicating higher organic pigment concentrations, while lighter backgrounds show attenuated or minimal pigment signals.

**Figure 4.**
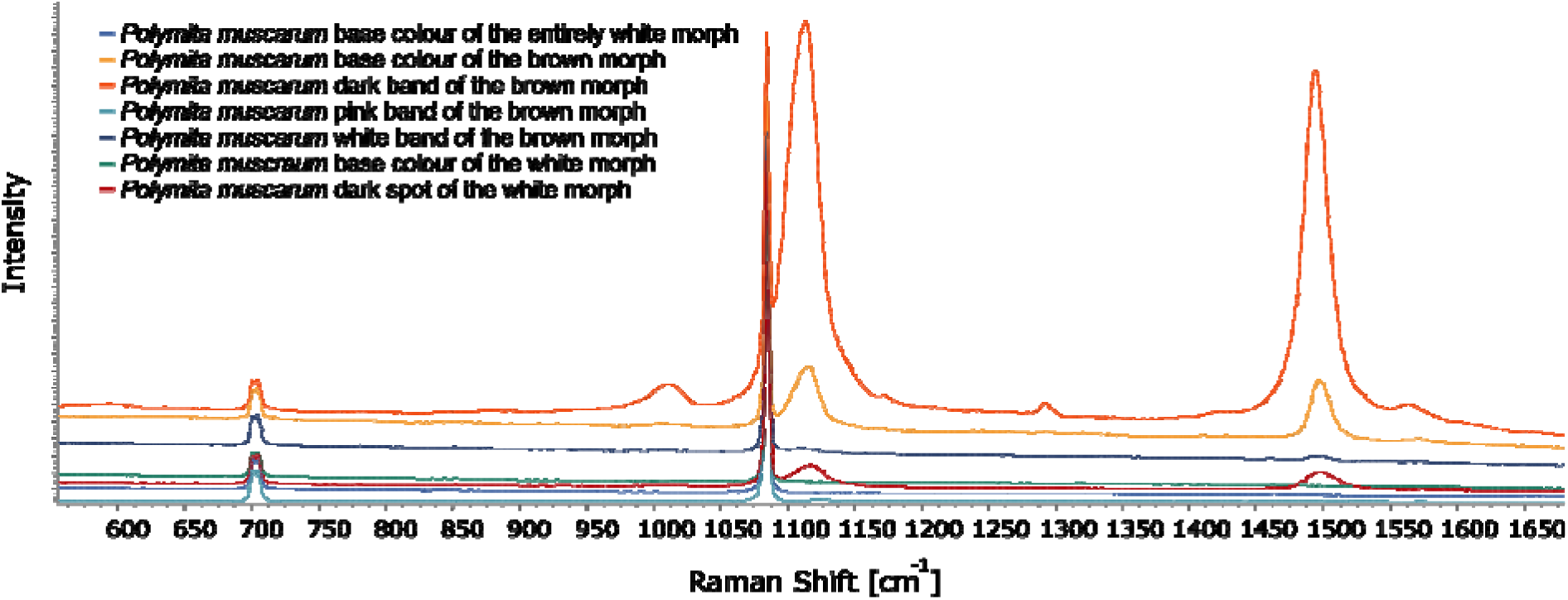
Raman spectra obtained from the base colour, bands and spots in shell fragments of *Polymita muscarum*.

### Scanning electron microscopy (SEM) of shell ultrastructure and light transmission

The shell of *Polymita picta* and *P. muscarum* exhibits a crossed-lamellar architecture. Large first-order lamellae are composed of parallel second-order lamellae whose orientation alternates between adjacent lamellar sets, producing the characteristic crossed pattern (Fig. 5A). At higher magnification, these lamellae consist of tightly packed elongate aragonite units, likely corresponding to third-order lamellae (Fig. 5B). Where preserved, the periostracum exhibited a characteristic honeycomb-like surface structure (Fig. 5C). In other cases, it appeared eroded, forming smoother surface patches, and in several fragments it was partly or completely absent (see Table 1).

**Figure 5.**
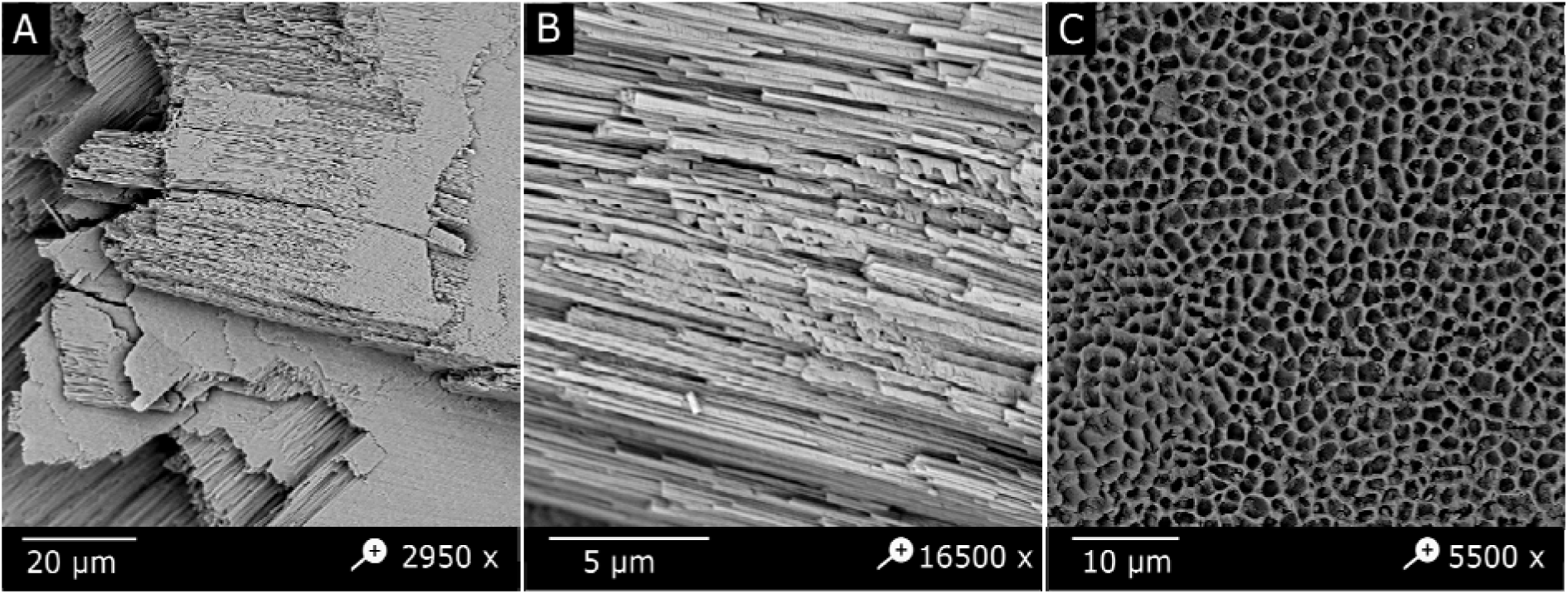
Details of the crossed-lamellar microstructure and the periostracum in the shell of *Polymita picta* (base colour red). A: Crossed-lamellar architecture showing the characteristic ∼60° insertion angle between adjacent sets of second-order lamellae. B: Higher magnification of the lath-like second-order lamellae. C: Honeycomb-like surface organization of the shell protein matrix in the outer surface of the periostracum.

**Table 1.** Measurements of shell layers (mean ± SD) in shell fragments of different colour morphs of *Polymita picta*. NA = layer absent.

| Morph | Periostracum ($\mu\text{m}$) | Outermost crossed-lamellar layer ($\mu\text{m}$) | Transverse crossed-lamellar ($\mu\text{m}$) | Co-marginal ($\mu\text{m}$) | Transverse crossed-lamellar ($\mu\text{m}$) | Co-marginal ($\mu\text{m}$) |
| --- | --- | --- | --- | --- | --- | --- |
| Black | 1.09 ± 0.22 | 28.43 ± 1.26 | 72.86 ± 1.38 | 23.58 ± 1.89 | NA | NA |
| Red | 2.44 ± 0.83 | 44.60 ± 3.06 | 63.67 ± 1.21 | 84.49 ± 1.39 | 177.95 ± 3.08 | |
| Green | NA | 37.11 ± 2.42 | 71.04 ± 3.31 | 60.72 ± 1.11 | 87.50 ± 2.77 | 16.58 ± 1.36 |
| Brown | 1.82 ± 0.57 | 23.82 ± 2.61 | 33.00 ± 4.50 | 54.63 ± 1.04 | NA | NA |
| Yellow | 1.87 ± 0.40 | 44.81 ± 0.70 | 81.76 ± 1.62 | 27.99 ± 2.12 | NA | NA |
| White | NA | NA | 75.67 ± 2.45 | 104.04 ± 2.92 | 135.67 ± 6.80 | 190.21 ± 5.31 |

First- and second-order lamellae are well delineated and the second-order ones consistently intersect at a characteristic angle of ∼60°. Local surface relief can occasionally impart a simple prismatic appearance, but the hierarchical arrangement is diagnostic of the crossed-lamellar type. On co-marginal fracture (e.g., fracture parallel to growth lines), the layering from the interior inwards is consistent in both species (Fig. 6).

**Figure 6.**
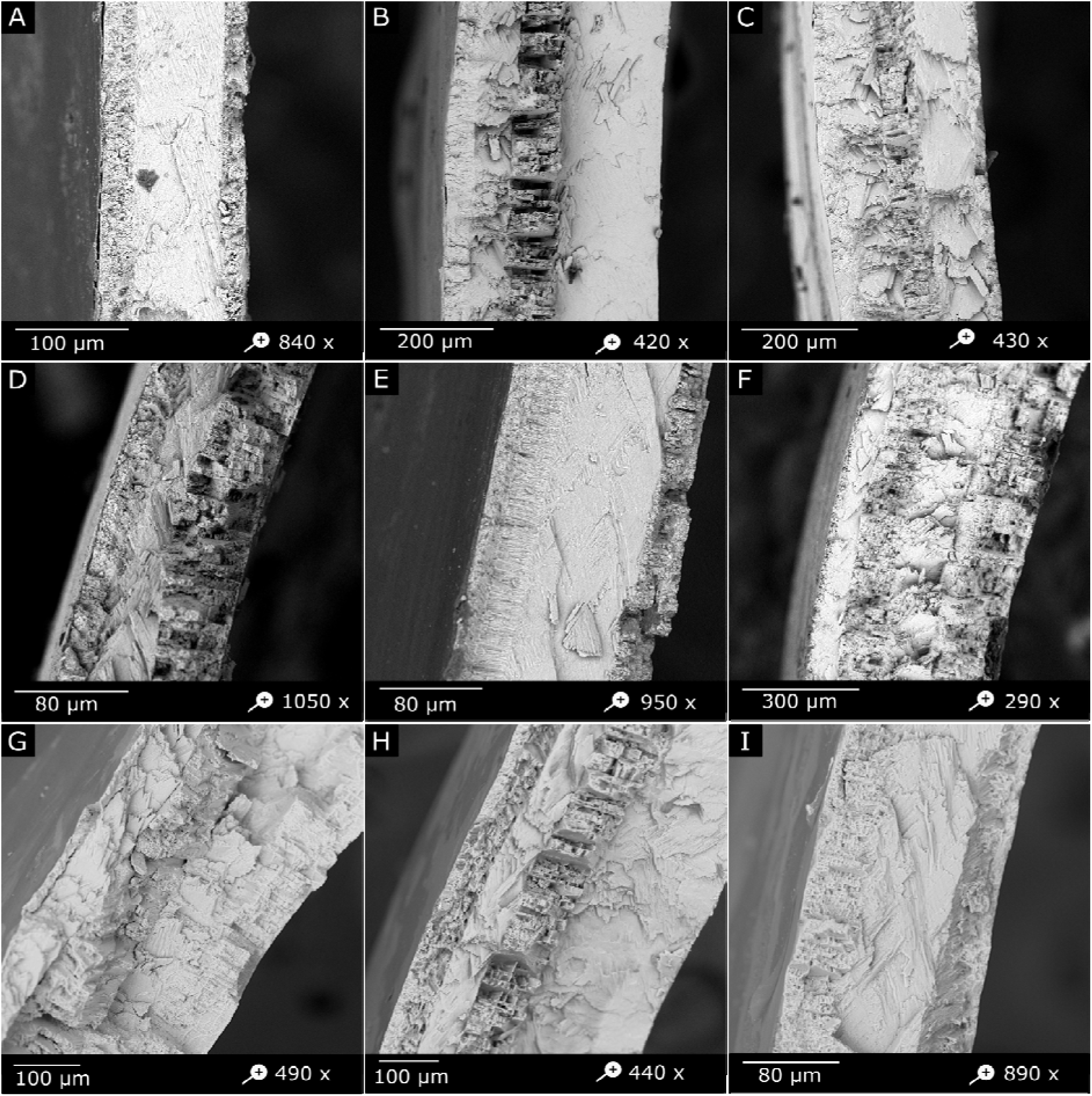
SEM images of co-marginal fractures in shells of *Polymita picta* and *P. muscarum*. The periostracum (where present) and the architecture of the shell layers are shown for different base colour morphs. A: *P. picta,* black morph. B: *P. picta,* red morph. C: *P. picta,* green morph. D: *P. picta,* brown morph. E: *P. picta,* yellow morph. F: *P. picta,* white morph. G: *P. muscarum,* brown morph. H: *P. muscarum,* white morph. I: *P. muscarum,* entirely white morph.

Immediately beneath the periostracum (where present; see Tables 1 and 2), there is an outer crossed-lamellar layer that inserts obliquely (∼45°) relative to the periostracal plane. Within this layer, one set of second-order lamellae is oriented approximately perpendicular to the shell surface, while a complementary set intersects it at an angle of ∼60°.

**Table 2.** Measurements of shell layers (mean ± SD) in shell fragments of different colour morphs of *Polymita muscarum*. NA = layer absent.

| Morph | Periostracum ($\mu\text{m}$) | Outermost crossed-lamellar layer ($\mu\text{m}$) | Transverse crossed-lamellar ($\mu\text{m}$) | Co-marginal ($\mu\text{m}$) | Transverse crossed-lamellar ($\mu\text{m}$) | Co-marginal ($\mu\text{m}$) |
| --- | --- | --- | --- | --- | --- | --- |
| Brown | NA | 17.87 ± 3.21 | 80.16 ± 19.59 | 79.22 ± 4.14 | 102.10 ± 6.86 | NA |
| White | NA | 61.13 ± 4.85 | 64.77 ± 2.59 | 89.74 ± 2.80 | 210.71 ± 1.82 | NA |
| Entirely white | NA | 49.09 ± 0.90 | 106.56 ± 1.46 | 8.90 ± 0.69 | NA | NA |

Below this outer layer, additional crossed-lamellar layers are arranged in succession, differing in their orientation relative to the shell growth direction. In particular, layers with lamellae arranged transversely alternate with layers in which the lamellae follow a co-marginal orientation. This alternating organization is repeated toward the inner shell (hypostracum), forming a stacked sequence of structurally distinct layers. The number of such layers and their individual thicknesses vary among shell fragments (see Table 1-2). Quantitative measurements supporting these observations are provided in Supplementary Data 3: https://doi.org/10.5281/zenodo.17343101. and are intended to support figure interpretation rather than species-level comparisons.

At the shell surface there may be spots that form punctate areas, which sometimes are partially covered by the periostracum. Intentional fracturing of shell fragments near such spots exposes their atypical architecture. In general, the spots occur in shell areas where the crystalline structure is less ordered than in the surrounding crossed-lamellar layer and which show an apparently higher porosity (e.g., larger inter-prismatic spacing) (Figs 7-8). Within spot areas, proteinaceous matrix elements are conspicuous, and the spot bases are bounded by a distinct organic layer. EDS signals consistent with nitrogen reached up to 22.83 % (Supplementary Data 4: http://doi.org/10.5281/zenodo.17343183), consistent with a potential protein-rich phase. In one individual of *P. muscarum* (brown morph), three spot areas showed average “depths” of 17.15 ± 0.27, 10.43 ± 0.16 and 6.60 ± 0.04 µm (Supplementary Data 3: https://doi.org/10.5281/zenodo.17343101).

**Figure 7.**
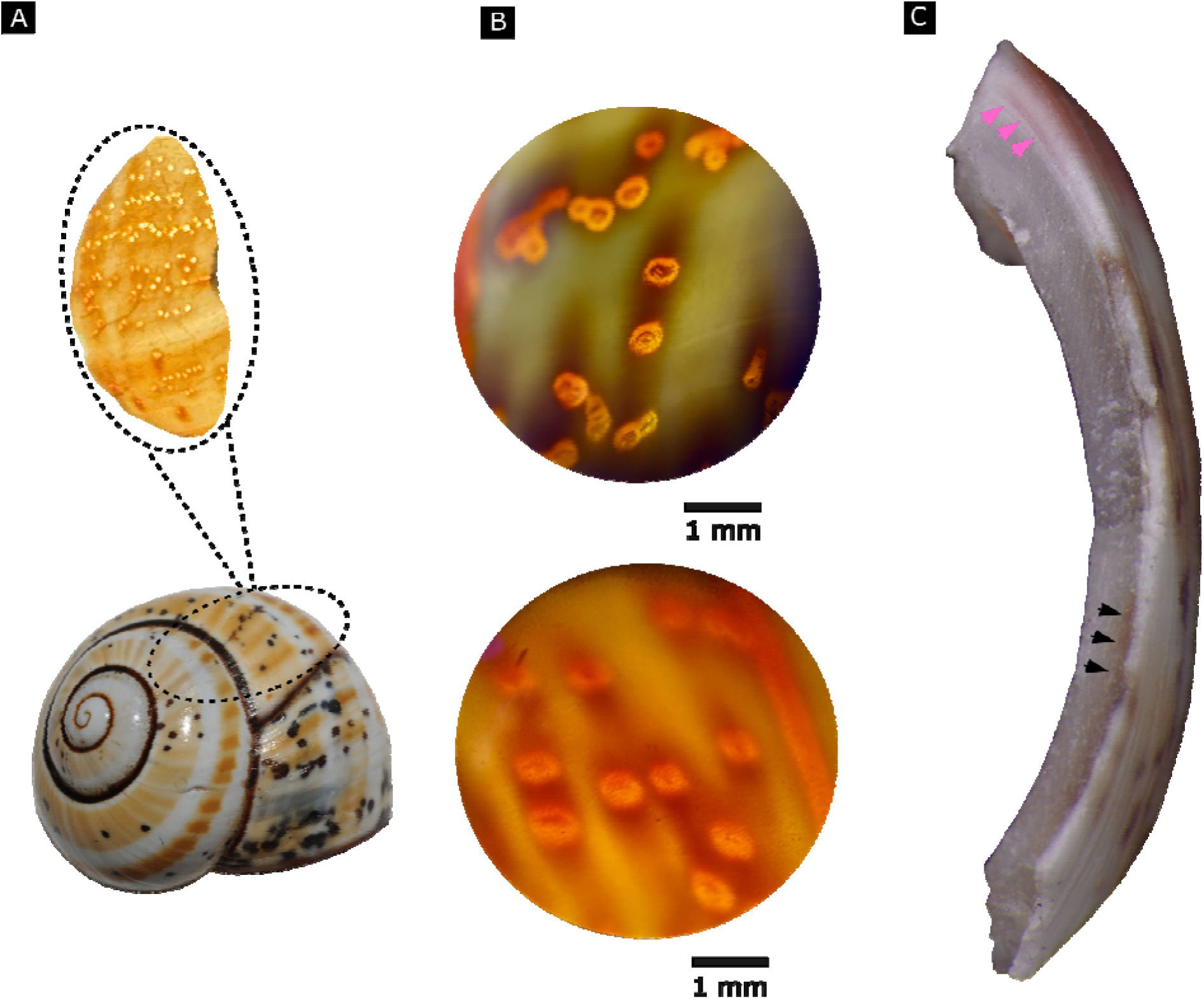
A: Light-transmitting spots in a punctate area in the shell of *Polymita muscarum* (brown morph), transilluminated along the direction of natural light incidence. B: Light microscopic detail (40x) of the light-transmitting spots viewed from outside (upper image) and inside (lower image) the shell. C: Microscopic (56x) transverse view of a *P. muscarum* shell fragment. Black arrows indicate a light-transmitting spot area and an underlying brown band; pink arrows indicate pink pigment deposition layers.

**Figure 8.**
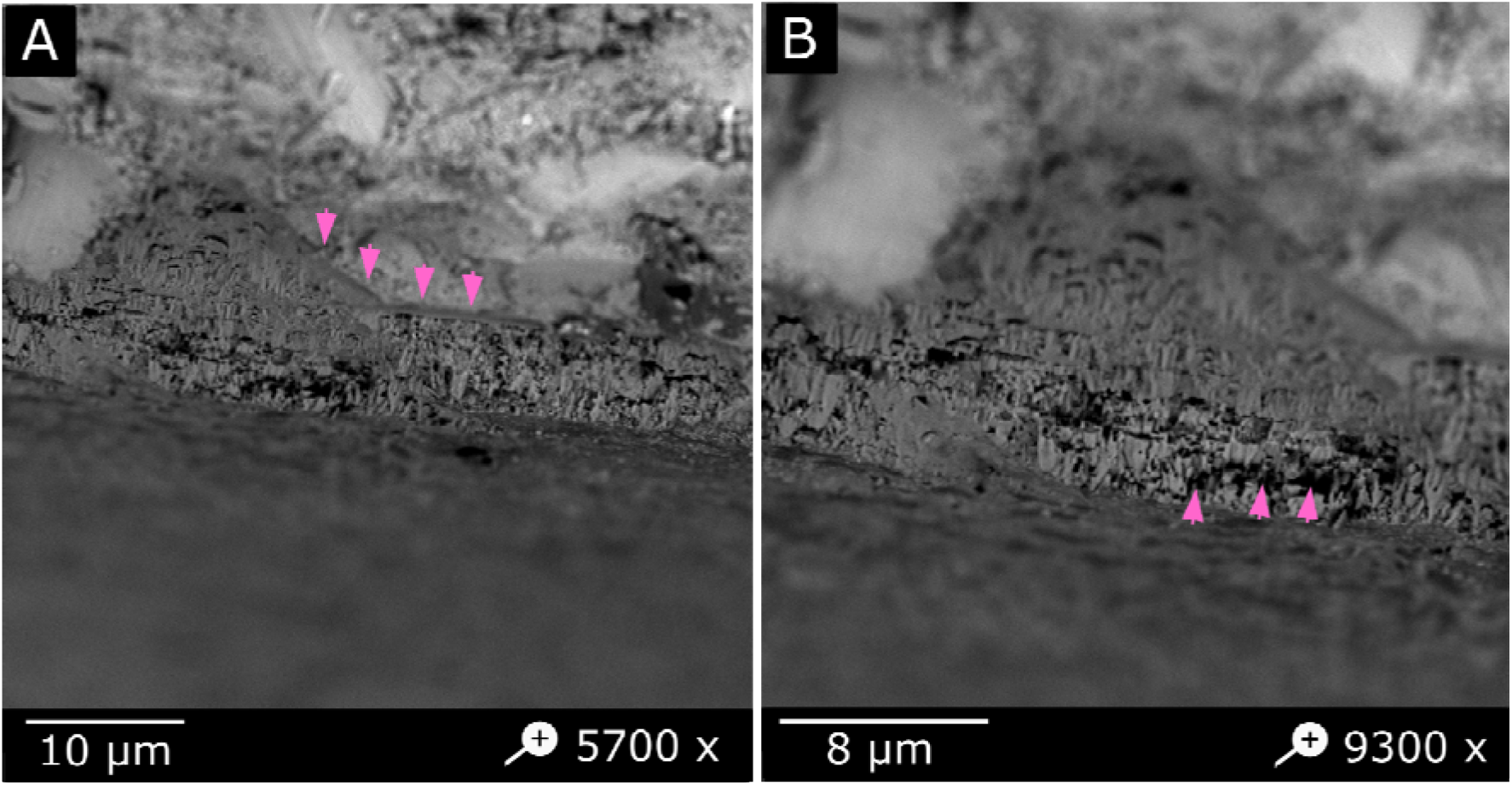
A-B: SEM images showing a lateral view of a light-transmitting shell spot in *Polymita muscarum* (brown morph). A: The darker microstructural domain of the spot is bordered by basal organic layer (marked by the arrows) and contrasts with the lighter surrounding shell mineral. B: Higher magnification of the central spot area partially covered by the periostracum. The increased intercrystalline spacing is indicated by arrows.

Under LED transillumination, shell spot areas act as a light-transmitting domains, that allow substantially more radiation to pass than the adjacent shell layers. This effect was consistent across shells, at least for the LED source used (Fig. 9). Shell areas with light-transmitting spots are common in *P. muscarum*, but rare in *P. picta*. In *P. muscarum*, the spots occur along most of the spire, extending nearly to the protoconch. Despite their prevalence, the spots do not seem to be distributed on the shell according to obvious spatial patterns.

**Figure 9.**
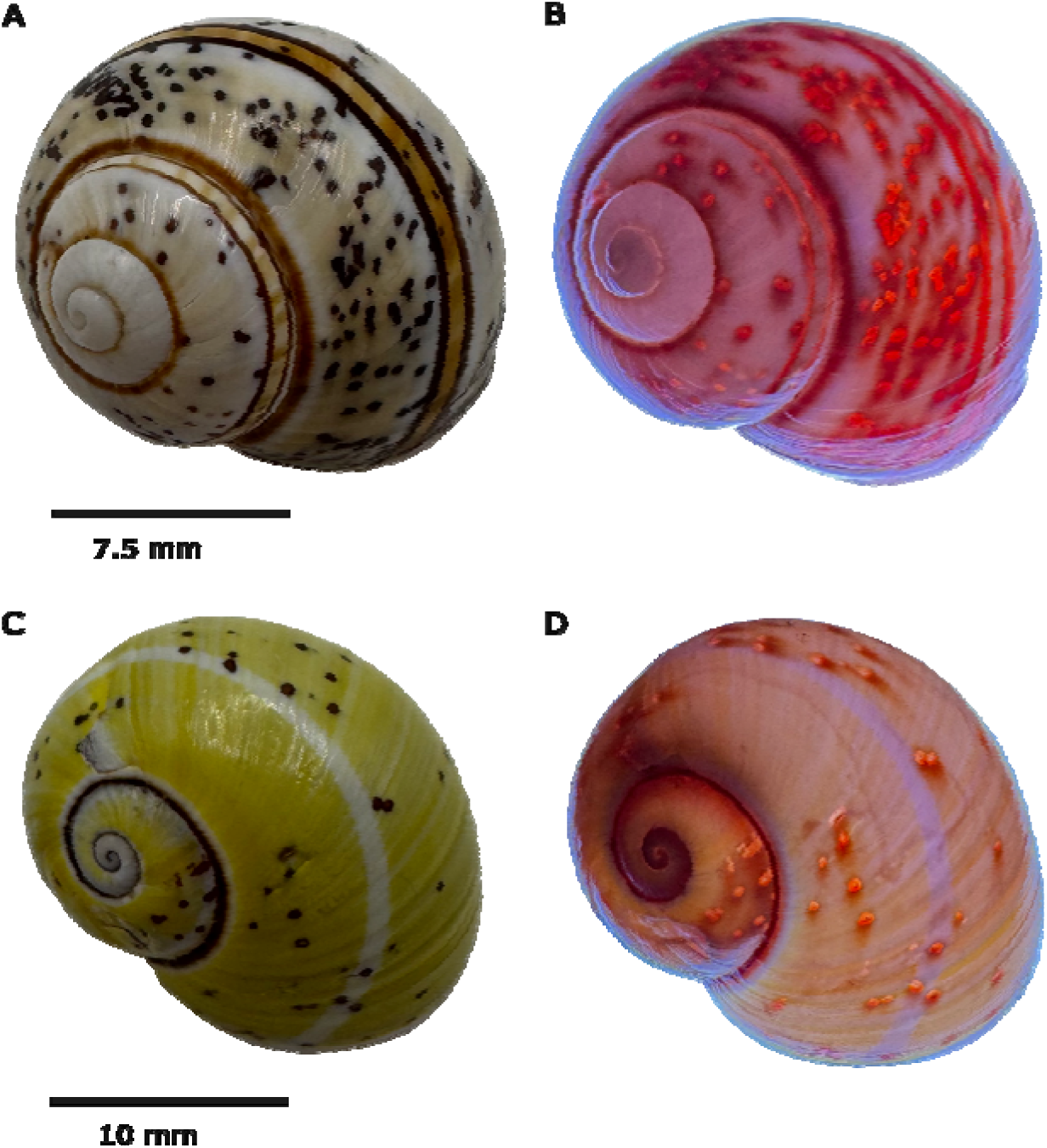
Light-transmitting spots in shells of *Polymita muscarum* (A, B) and *P. picta* (C, D) revealed by transillumination. A, C: under reflected light. B, D: transilluminated with white LED light to reveal light transmission.

## DISCUSSION

To better understand the possible ecological roles of the colour polymorphism in the shells of *Polymita picta* and *P. muscarum*, we explored the shell pigments and crystalline ultrastructure of different shell colour morphs of both species.

### Presence and concentration of total melanin in shell colour morphs of *Polymita picta* and *P. muscarum*

We identified and quantified total melanin in shell fragments of both species by measuring UV absorbance at 215 nm and comparing these values to a standard calibration curve based on *Sepia officinalis* melanin. Absorbance at this wavelength must be interpreted with caution, as several organic compounds can contribute to signals in the 200-220 nm range (Scopes, 1974; Jiang et al., 2010; Soltani et al., 2019). To minimize such interference, shell material was subjected to prolonged acid hydrolysis and thermal treatment following Hao et al. (2015), which removes most mineral and soluble organic components. This procedure yields an insoluble pigment fraction enriched in melanin-like compounds, allowing comparative quantification based on UV absorbance. The consistent detection of melanin-like spectra across all samples supports the presence of melanin in the analysed shell morphs, although the values should be interpreted as melanin-equivalent estimates rather than absolute concentrations. Quantification was performed on pooled samples and should therefore be interpreted cautiously, as no replication at the individual level was available. In addition, a secondary absorption feature around ∼270–280 nm was observed in darker morphs (Fig. 2I). This feature is consistent with previously reported spectra of melanin oxidation products such as pyrrole-2,3-dicarboxylic acid (PDCA) (Affenzeller et al., 2019a), although its identification here remains tentative.

### Raman spectroscopy

The Raman spectroscopy data showed that the shells of both *Polymita picta* and *P. muscarum* mainly consist of aragonite, consistent with most other pulmonate shells (Alves et al., 2023; Li et al., 2025). Additional Raman spectroscopy signals correspond to the canonical carotenoid resonance Raman triplet, with expected shifts associated with conjugation length, protein interactions and aggregation state (Withnall et al., 2003; Macernis et al., 2014; Udensi et al., 2022; Kolašinac et al., 2025).

Eumelanin typically produces broad D/G-like envelopes around ∼1380 and ∼1580 cm^−1^, but at 785 nm excitation its detection is hampered by strong fluorescence backgrounds and low signal-to-noise ratios (Ferrari et al., 2000; Huang et al., 2004). The absence of clear melanin peaks therefore does not exclude its presence, as supported by independent UV (215 nm) based estimates. A more definitive characterization of eumelanin versus pheomelanin would require alkaline H_2_O_2_ oxidation followed by HPLC of diagnostic markers, i.e. pyrrole-2,3,5-tricarboxylic acid (PTCA) and pyrrole-2,3-dicarboxylic acid (PDCA) for eumelanin and thiazole-2,4,5-tricarboxylic acid (TTCA) and pyrrole-4,5-tricarboxylic acid (TDCA) for pheomelanin (d’Ischia et al., 2013).

Our findings in carotenoid-related Raman spectra signals are consistent with those reported in neritid shells (Komura et al., 2018), where carotenoids accumulate progressively from inner layers, while melanin is restricted to a narrow surface area. In *Polymita*, the presence of pronounced polyene bands (Udensi et al., 2022) in (1) the dark colour bands of both species, (2) the black and red base colours of *P. picta,* and (3) the base brown colour and spots of *P. muscarum*, supports a mixed-pigment model in which carotenoids and melanins jointly contribute to the observed colour variation. This interpretation is consistent with previously reported continuous variation in shell luminance across morphs (Gordillo-Pérez et al., 2025), suggesting that colour continuum can be explained by shifts in the relative contribution of co-occurring pigments.

### Shell structure and light-transmitting shell spots

The SEM images showed that both species have a crossed-lamellar shell microstructure comprising first-, second- and third-order lamellae, which is the most frequent arrangement in gastropods (de Paula and Silveira, 2009). The most evident structural feature is the intersection between two sets of second-order lamellae, which meet at characteristic angles of approximately ∼60°. In gastropods, this hierarchical architecture provides high fracture resistance through crack deflection across multiple structural length scales (de Paula and Silveira, 2009; Marin et al., 2012; Rodríguez-Navarro et al., 2012). The alternation of crossed-lamellar layers with co-marginal layers, repeating inward toward the hypostracum, is consistent with established models of crossed-lamellar shell organization and with previous reports indicating species-specific variation in layer thickness (Checa, 2018). In *P. picta*, however, the thickness and relative proportions of these topologically equivalent shell layers appear to vary among the examined colour morphs. Whether this variation reflects true differences associated with shell colour morphs, or instead results from individual or ontogenetic variability requires further investigation.

The periostracum, which did not always cover the entire shell surface, likely reflecting natural wear or post-collection degradation, showed a honeycomb-like organic microstructure. Such structures are consistent with previously described periostracal and inter crossed-lamellar matrices that influence nucleation and orientation of aragonite and may locally affect pigment incorporation within the shell (Chateigner et al., 2000; de Paula and Silveira, 2009; Nakayama et al., 2013). However, not in every shell fragment of *Polymita*, the periostracum presented the honeycomb-grid organization, for in some cases the outer layer was smooth or partially, if not totally, absent. This aspect warrants further investigation to clarify its potential role in shell colour.

At the shell surface of *Polymita muscarum*, and in some shells of *P. picta*, spots occur in porous areas enriched in organic (protein-rich) components that interact differently with light. SEM reveals punctate surfaces which, upon fracture adjacent to these spots, show reduced crystalline order and increased porosity (i.e. larger inter-prismatic spacing) relative to the surrounding crossed-lamellar layer. These regions also display a conspicuous proteinaceous phase and a basal organic layer, consistent with localized organic enrichment. Elemental analyses (EDS) of light elements (e.g. C, N, O) in heterogeneous biominerals are inherently semi-quantitative due to analytical limitations (e.g. absorption of low-energy X-rays and sensitivity to sample geometry; Gazulla et al., 2013; Newbury et al., 2015). Within this constraint, the relatively elevated nitrogen signal observed in spot regions supports a higher organic content compared to the surrounding shell material.

### Cryptotransmissive domains

Taken together, the observed features of the shell structure explain the transillumination behaviour of isolated spots within the shells. Although these spots contain pigment and appear dark in most morphs, the lower density of the mineral matrix renders them less opaque to transmitted light. These areas function as “windows” (or pores for the individual shell spots) that are abundant in *P. muscarum* but comparatively sparse in *P. picta*. To our knowledge, such light-transmitting shell structures have not been described in this form in gastropods.

It is important to distinguish these structures from shell pores and light-sensitive systems described in other molluscs, such as shell eyes in polyplacophorans (Lee et al., 2011; Li and Ortiz, 2013; Li et al., 2015; McCoy et al., 2024), which are specialized photoreceptive organs associated with sensory cells. In contrast, the shell spots described here appear to be passive structural features that allow the transmission of light, rather than photoreceptive systems. Shell pore and channel systems have been described in other gastropods, particularly in caenogastropod lineages such as Cyclophoroidea, where microtunnels and sutural tube systems form complex shell-integrated structures involved in gas exchange (Páll-Gergely et al., 2016, Páll-Gergely, 2023, 2025). These consist of networks of narrow tunnels penetrating the shell layers and connecting internal and external environments. However, the *Polymita* shell spots do not appear to be directly comparable to these systems in terms of morphology, organization, or inferred function. Channel-like or porous structures have also been reported in other molluscan groups, including bivalves, where they may be associated with structural, physiological, or biomineralization-related roles (e.g. Batten, 1984; Okada et al., 2019). Taken together, these observations indicate that shell-associated channel systems occur across Mollusca and may fulfil diverse functions. Within this broader context, the light-transmitting shell spots observed in Polymita are best interpreted as a distinct type of shell feature, the homology of which with other molluscan channel systems remains uncertain. Rather than representing a direct analogue of previously described systems, they may constitute a previously unrecognized form of shell tunnel structure. The irregular spatial distribution of these spots across the shell surface, extending up to the protoconch, further suggests that their formation does not follow a regular periodic morphogenetic pattern.

Their capacity to transmit light suggests that these *Polymita* cryptotransmissive domains may act as “windows” to provide light cues to animals that are withdrawn into the shell. This interpretation is based on (i) the presence of discrete shell areas that allow light transmission, (ii) their distribution across the shell surface, and (iii) the observed association between spot-rich regions and increased shell translucence. As such, they could potentially trigger or modulate a snail’s behaviour and/or physiology in relation to, for example, thermoregulation. However, this sensory interpretation remains circumstantial. Demonstrating that the light-transmitting shell spots in *Polymita* effectively function as a bona fide light-sensory system would require targeted behavioural assays under controlled illumination, mapping of light sensitivity in mantle tissues, and exclusion of alternative structural-optical explanations. Optical imaging studies in land snails suggest that translucence depends strongly on both shell pigmentation and microstructure (Savazzi and Sasaki, 2013), which is consistent with our observation that in *Polymita* shells, surface areas with many spots are particularly light-transmissive.

### A tentative integrative interpretation of colour polymorphism in *Polymita*

The joint presence of melanin and carotenoids in *Polymita* shells provides the basis for their shell colour polymorphism. Both pigment classes (and potentially their oxidation products) may buffer light radiation and heat stress by altering light absorbance and dissipation (Emberton et al., 1963; Schweizer et al., 2019; Gordillo-Pérez et al., 2025). This view inspires us to speculate that shell colour polymorphism in *Polymita* may have a thermoregulatory function, based on the spatially heterogeneous distribution of pigments within the shell. At the intra-shell level, pigments are unevenly distributed across bands, spots, and background areas, generating local differences in optical properties that could help modulate heat absorption and stabilize body temperature under fluctuating insolation. This organization creates a micro-framework for an organic matrix that regulates crystalline orientation and pigment distribution, thereby influencing optical behaviour via local refractive-index contrast and scattering (Rodríguez-Navarro et al., 2012; Osuna-Mascaró et al., 2014; Bonnard et al., 2020; Agbaje et al., 2021). In this view, the crossed-lamellar shell architecture is not a passive backdrop but instead provides a hierarchical interface through its variable inter-lamellar spacing. *Polymita* colour polymorphism is not only aesthetic. The organisation of pigments and minerals, the ecological interactions of these structural elements, and their origin and evolution likely holds the key to potential thermoresistance and climate change adaptation.

## Notes

### Competing Interest Statement

The authors have declared no competing interest.

### Summary of Updates

Aspects of the abstract, the calibration curve and calculations, Figure 9, and the discussion have been revised in agreement with the authors.

